# Characterization of the *Cannabis sativa* glandular trichome proteome

**DOI:** 10.1101/2020.11.09.373910

**Authors:** Lee J. Conneely, Ramil Mauleon, Jos Mieog, Bronwyn J. Barkla, Tobias Kretzschmar

## Abstract

*Cannabis sativa* has been cultivated since antiquity as a source of fibre, food and medicine. The recent resurgence of *Cannabis* as a cash crop is mainly driven by the medicinal and therapeutic properties of its resin, which contains compounds that interact with the human endocannabinoid system. Compared to other medicinal crops of similar value, however, little is known about the biology of *C. sativa*. Glandular trichomes are small hair-like projections made up of stalk and head tissue and are responsible for the production of the resin in *C. sativa.* Trichome productivity, as determined by *Cannabis sativa* resin yield and composition, is only beginning to be understood at the molecular level. In this study the proteomes of glandular trichome stalks and heads, were investigated and compared to the proteome of the whole flower tissue, to help elucidate *Cannabis sativa* glandular trichome biochemistry. The data suggested that the floral tissue acts as a major source of carbon and energy to the glandular trichome head sink tissue, supplying sugars which drive secondary metabolite biosynthesis in the glandular trichome head; the location of the secretory cells. The trichome stalk seems to play only a limited role in secondary metabolism and acts as both source and sink.

## Introduction

*Cannabis sativa* is an annual, predominatly dioecious [1], monotypic species of the Cannabaceae in the order *Rosales* [2]. Its center of diversity is in central Asia, which was also proposed as its center of origin [3]. The use of *Cannabis sativa* can be traced back for millenia through both written and genetic evidence with the earliest plant remains found in burial chambers in Yanghai, China, dated to approximately 2,500 years ago [4]. *Cannabis sativa* has many uses. The seeds are a source of oil and protein rich nutrition [5], the fibres are raw material for the production of clothing and rope [6], and the resin has proven therapeutic properties with medicinal, religious, cultural, and recreational applications [7]. Due to the narcotic effect of some cultivars of *Cannabis sativa* the plant was declared a controlled drug or prohibited substance in the first half of the 20^th^ century in most jurisdictions and, under the United Nations single convention on narcotic drugs [8], remains highly regulated in most countries.

Cultivation of *Cannabis sativa* as a medicinal crop is increasing rapidly and is driving a research renaissance in the plant science community [9]. This is in part due to relaxation of *Cannabis* legislation in several developed countries, which has fuelled an unprecedented growth of legal *Cannabis* and derived products, creating multi-billion dollar markets in the US alone [10, 11].

Glandular trichomes are specialised multicellular epidermal protrusions that function as miniature secondary metabolite producing factories [12]. They are found in approximately one third of all angiosperms, where they fulfill a variety of roles in plant-environment interactions [13]. The production of distinct secondary metabolites in glandular trichomes is often species specific [14, 15]. While terpenoids are arguably the largest class of secondary metabolites produced by glandular trichomes, other common compound classes include phenylpropanoids [16], acyl sugars [17], flavonoids [18] and methylketones [19]. Cannabinoids, a class of terpenophenolic compounds [20], that interact with the human endocannabinoid system, are exclusively found in *Cannabis sativa* [21].

Capitate stalked glandular trichomes are the most abundant trichome type found on *Cannabis sativa* [22]. Cannabinoids and terpenoids are synthesised by a cluster of secretory cells located at the base of the glandular trichome head, referred to as disc cells. Secondary metabolites accumulate in the sub-cuticular cavities located between the disc cells and the cuticle of the glandular trichome head. The trichome head sits on top of a multicellular stalk connected by stipe cells [23]. The stalk is continuous with the epidermis of the flower tissue, and the walls are notably cutinised. It has been suggested that cutinisation of the stalk cell wall is to prevent leakage of resin produced by the gland head into the apoplastic space [24].

The capitate stalked glandular trichomes are particularly abundant on the female reproductive organs – floral leaves and bracts - of *Cannabis sativa* [25]. This is likely an adaptation to protect developing seed from herbivory, thus enhancing chances of reproductive success [26]. The exceptionally high productivity of *Cannabis sativa* glandular trichomes with respect to cannabinoid production, however, is likely a consequence of artifical selection and targeted breeding [27].

The two main cannabinoids, cannabidiol (CBD) and tetrahydrocannabinol (THC) are well documented for their therapeutic properties [28, 29], which is largely mediated through interaction with the human endocannabinoid system [30]. CBD has been shown to have anti-inflammatory properties and is capable of increasing the pro-apoptotic abilities of human immune cells [31], whereas THC is the main psychoactive compound [32]. In addtion to CBD and THC over 100 additional cannabinoids have been detected using chromatography and mass-spectrometry based techniques [33, 34]. They are typically referred to as minor cannabinoids and include cannabigerolic acid (CBGA), cannabichromenic acid (CBCA), cannabinolic acid (CBNA), and cannabicyclolic acid (CBLA), with several showing potential to alleviate different disease states [35] [36].

A key enzyme in the biosynthesis of cannabinoid precursor molecules is polyketide synthase III, a tetraketide synthase, that catalyses the condensation of three malonyl Co-A with one molecule of hexanoyl Co-A to produce polyketide intermediates for olivetolic acid cyclase (OAC) to produce olivetolic acid (OA) [37]. Cannabigerolic acid (CBGA) is synthesised through conjugation of OA and the isoprenoid geranyl pyrophosphate (GPP) via the enzymatic activity of a prenyltransferase, *Cannabis sativa* prenyltransferase 4 (CsPT4), sometimes referred to as cannabigerolic acid synthase [38]. CBGA is a substrate for tetrahydrocannabinolic acid synthase (THCAS), cannabidiolic acid synthase (CBDAS) and cannabichromenic acid synthase (CBCAS), producing tetrahydrocannabinolic acid (THCA), cannabidiolic acid (CBDA), and cannabichromenic acid (CBCA) respectively. Prenyltransferases such as CsPT4 demonstrate flexibility in substrate specificity and are known to accept derivatives of OA with variable alkyl side chain lengths [39].

Another class of cannabinoids known as cannaflavins are produced through the enzymatic activity of *C. sativa* prenyltransferase 3 (CsPT3). CsPT3 catalyses the covalent linkage of GPP or dimethylallyl pyrophosphate (DMAPP) with the methylated flavone chrysoeriol to produce cannaflavins A and B [40]. While the biosynthetic pathways of the major cannbinoids have been mapped to a large degree, little is known about the drivers of trichome productivity that enable these compounds to be accumulated in such abundance.

Driven by the economic value of trichome-specific secondary metabolites, which have found use as pharmaceuticals, food flavourings, and insecticides, there has been extensive research conducted into the biology of glandular trichomes. A main focus has been on understanding the proteins, pathways, and regulatory mechanisms governing the biosynthesis of secondary metabolites [12, 41]. In the long term these activities have been proposed to aid in developing strategies to bioengineer systems for increased production of commercially valuable secondary metabolites [13, 42–45]. In this context the proteomes of trichomes across a range of different species including *Solanum lycopersicum, Artemisia annua, Ocimum basilicum L., Olea europaea* and *Nicotiana tabacum* [46–50] have been analysed. Investigation of the proteins found in *Artemesia annua* glandular trichomes, by means of a comparative proteomics approach revealed that proteins involved in the electron transport chain, isoprenoid biosynthetic proteins, translation, and proteolysis [47] were more abundant in the trichomes as compared to leaf tissue. A shotgun proteomic analysis of glandular trichomes isolated from *Solanum lycopersicum* revealed all steps of the methylerythritol 4-phosphate (MEP) pathway, as well as proteins involved in rutin biosynthesis, and terpene biosynthesis [48] to be over-represented. Furthermore, glandular trichomes were associated with high energy production and protein turnover rates.

Proteomic analysis of *Cannabis sativa* was pioneered by Raharjo *et al.* in 2004 [51]. Using two-dimensional gel electrophoresis and mass spectrometry, 300 proteins were identified from the late-stage flowers of *Cannabis sativa.* A further 100 proteins were identified from “gland extracts”, likely a mix of epidermal tissues enriched for glandular trichomes. Proteins identified included zinc-finger type proteins, F-box family of proteins, and a range of secondary metabolite synthases. Notably lacking were proteins involved in the biosynthesis of cannabinoids. A recent report on the proteome of *Cannabis sativa* glandular trichomes focused on the development of optimal protein extraction methods from apical flower buds and isolated glandular trichomes, however, subsequent shotgun proteomics on the isolated samples only identified 160 proteins in *C. sativa* mature apical buds and glandular trichomes [52]. The approach taken also made it difficult to distinguish proteins specific to glandular trichomes from those proteins specific to the apical buds. A number of secondary metabolite biosynthetic proteins, including those in the cannabinoid pathway, were identified. The combined proteome of the apical bud and glandular trichome included terpene synthases, members of the plastidal MEP pathway, and three members of the cannabinoid biosynthetic pathway, olivetolic acid cyclase (OAC), tetrahydrocannabinolic acid synthase (THCAS), and cannabidiolic acid synthase (CBDAS).

In this study 1240 proteins were identified in glandular trichome head protein isolates, 396 proteins from glandular trichome stalk protein isolates, and 1682 proteins in *C. sativa* late-stage flowers, using quantitative time of flight mass spectrometry (QTOF-MS/MS) and matching assembled peptides to the *Cannabis sativa* cs10 reference proteome (NCBI). Analysis of relative protein abundance of glandular trichome heads, stalks, and late-stage flowers was used to idenitfy proteins that were shared between tissues, those that were tissue specific, and proteins which showed significant differential levels of abundance. Comparison of glandular trichome protein head isolates to the *C. sativa* late-stage flower protein isolates showed the extent to which glandular trichome heads are governed by secondary metabolite biosynthesis, secondary metabolite transport, and other processes such as carbon refixation.

## Methods

### Plant growth conditions

Seeds of *Cannabis sativa* cultivar hemp No.8 were germinated on paper towels moistened with 0.01% gibberellic acid in water. After 7 days, seedlings were transferred to 1L pots containing a mix of 1:1:1 vermiculite, perlite, peat moss, supplemented with 1 g/L dolomite. Seedlings were grown under conditions promoting vegetative growth for five weeks in a Sanyo growth cabinet (18hr light, light level 5). Plants were watered every three days with CANNA veg (CANNA) (40mL A + 40mL B/ 10L water). For reproductive growth, plants were transferred to a growth room and placed under a Vipar spectra grow light (LED model: R900) (900W)) under a 12:12 h light/dark cycle with both veg and bloom settings switched on. During flowering, plants were watered every three days with water and once per week with CANNA flor (40mL A + 40mL B/ 10L water) and grown for approximately eight weeks or until two thirds of stigmas were brown in colour, indicating floral maturity.

### Scanning electron microscopy and fluorescence microscopy

Scanning electron microscopy - late-stage *Cannabis sativa* flowers were harvested, and the calyx was removed and placed on an adhesive carbon strip mounted on a loading stage for imaging. A Hitachi TM4000 plus scanning electron microscope was used with the cold stage set to −30°C for sample mounting. Voltage was set to 10kV – mode three and standard (M) vacuum.

Fluorescence microscopy – calyx from late-stage *Cannabis sativa* flowers were harvested using a blade and mounted on a glass slide. Images were taken using a Nikon E600 fluorescent microscope at an excitation wavelength of 400-440 nm with Dichromatic filter (DM) set at 455 nm and Barrier filter (BA) set at 480 nm. Images were captured using an Olympus DP72 camera controlled through the Nikon Cellsense software.

### Isolation of glandular trichomes

Epidermal tissues were harvested from late-stage (week 7) *Cannabis sativa* flowers. One gram of late-stage flowers was flash frozen in liquid N_2_ and grated against a 425μm mesh sieve. Grated tissue was collected into a 150μm mesh sieve. Liquid N_2_ was poured over the 150μm mesh to facilitate passage of particles onto a steel Pyrex™ collecting dish. The sieved material was then collected using precooled waxed paper, and 3 mL rapidly transferred into a prechilled 15 mL falcon tube, followed by the addition of 3mL of prechilled mannitol buffer (0.2 M mannitol, 0.05M Tris-HCl, 0.02 M sucrose, 0.005M MgCl_2_, 0.01M KCl, 0.0005M K_2_HPO_4_, and 0.001 M EGTA).

The epidermal tissue extract was layered onto previously established discontinuous Percoll® density gradients consisting of a cushion of 80% Percoll® followed by 3mL of 60%, 45%, and 30% Percoll® in mannitol buffer. Fractionation of the epidermal tissue mixture was achieved by centrifugation of the discontinuous Percoll® density gradient at 400 *g* at 4 °C for 10 min [53]. Glandular trichome heads were recovered at the 0/30% Percoll® interface.

Glandular trichome stalks were recovered at the interface of the 30% and 45% Percoll layers. Fractions were collected from the gradient using a P1000 pipette and transferred into 2mL Eppendorf tubes. Samples were washed free of residual Percoll^®^ using 5 volumes Milli-Q water. Material was pelleted by centrifugation at 1000 *g* for 5 mins in a refrigerated benchtop centrifuge (make and model) and the supernatant was removed. The wash step was repeated three times. Samples were flash-frozen in liquid N_2_ and stored at −80 °C.

### Protein isolation

Total protein was isolated using approximately 100 mg of glandular trichome heads, 100 mg of glandular trichome stalks, or 1 g of flowers. Frozen samples were ground into a powder using a mortar and pestle. Powdered tissue was collected and placed into prechilled Eppendorf tubes, five volumes of (1:9) trichloroacetic acid (TCA)/ acetone [54] were added to each sample and mixed by inverting the tube several times. The suspension was filtered into a falcon tube using a 40 μm nylon mesh sieve. The filtrate was then stored overnight (O/N) @ −20 °C to allow for complete precipitation of protein. Samples were then centrifuged at 2000 *g* for 10 min in a tabletop refrigerated centrifuge, and the supernatant removed. Next, samples were washed with 5 mL of ice-cold acetone, and centrifuged at 2000 g for 10 min. The acetone wash step was repeated three times and the supernatant removed. Protein pellets were dried in a fume hood.

### Trypsin enzymatic digest

Protein pellets were resuspended (1mg/mL) in a buffer containing 2M urea and 50mM ammonium bicarbonate (pH 8) by vortexing and sonicated for 10 min. Tryptic digestion of protein samples was carried out by combining 20 μg of trypsin with 100 μL aliquots of each protein suspension. Tryptic digest reactions were carried out O/N @ 37 °C. Samples were lyophilized using a vacuum centrifuge. Lyophilised samples were dissolved in 100 μL 1% trichloroacetic acid (TCA)/Milli-Q water and transferred to glass sample vials.

### nanoHPLC Mass spectrometry

Protein samples were analysed on an Eksigent, Ekspert nano LC400 ultra HPLC coupled to a Triple time of flight (TOF) 6600 mass spectrometer (SCIEX) with a picoview nanoflow ion source. Five μL injections were run on a 75 μm x 150mm chromXP C18CL 3 μm column with a flow rate of 400 nL per minute and a column temperature of 45 °C. Solvent A consisted of 0.1% formic acid in water, and solvent B consisted of 0.1% formic acid in acetonitrile. Ionspray voltage was set to 2600V, de-clustering potential (DP) 80V, curtain gas flow 25, nebuliser gas 1 (GS1) 30, and interface heater at 150 °C. A linear gradient of 5-30% solvent B over 120 minutes at 400nL/minute flow rate, followed by a steeper gradient of 30% to 90% solvent B for 3 minutes, then 90% solvent B for 17 min was carried out for peptide elution. The mass spectrometer acquired 100ms full scan TOF-MS followed by up to 50 ms full scan product ion data in an information dependent acquisition manner. Full scan TOF-MS data was acquired over the range 350-1500 m/z and for product ion MS/MS 100-1500/ ions observed in the TOF-MS scan exceeding a threshold of 100 counts and a charge state of +2 to +5 were set to trigger the acquisition of product ion, MS/MS spectra of the resultant 50 most intense ions.

### Protein identification

Spectral data generated from nanoHPLC MS/MS was matched to peptides derived from protein sequences of the list of all proteins available for hemp in the Uniprot database as well as the list of predicted proteins derived from the *Cannabis sativa* cs10 project (https://www.ncbi.nlm.nih.gov/genome/gdv/browser/genome/?id=GCF_900626175.1) using the input file from ProteinPilot™ software version 5.0.1. The resulting MZidentifier file was exported to Scaffold4 software (Proteomesoftware) in order to view and quantitate the identified proteins found in our sample dataset.

### Protein data processing: normalisation and relative protein abundance

Normalization and relative protein abundance between samples was carried out using Scaffold4 software. Total spectra across all samples were normalized to allow for a semi-quantitative comparison of protein abundance. Total spectra were normalized using the normalized spectral abundance factor (NSAF) [55] whereby the assigned NSAF score for a protein is the number of spectra for a given protein in a sample divided by the length of that protein, further divided by the sum of all spectra of all proteins for a given sample. Total spectral counts of the aforementioned samples and their biological replicates were comma-separated variant (CSV) formatted and exported to PANDA view proteomics software [56]. For 0 value spectral counts, 0.001 was used to replace null values and the natural logarithm (Log_e_) was determined for all spectral counts for all samples. To compare the relative protein abundance between two sample types e.g. glandular trichome heads and stalks a t-test P<0.05 with Benjamini-Hochberg multiple test correction was implemented. The pairwise comparison of relative protein abundance of two sample groups e.g. glandular trichome heads and stalks could then be plotted on a volcano plot i.e. Log_2_ Fold change on the x-axis vs −Log_10_ P-value. Allowing estimation of the proteins that are overexpressed or exclusive to a given sample. Only those proteins present in all three biological replicates, with at least two unique peptides, and passing the significance threshold with a fold change of at least 2x (Log2(fold change) ≥ 1; ≤−1) were analysed further.

### Identification of metabolic pathways and other processes by MapMan

Metabolic pathways and various processes represented by the lists of proteins (from the previous step) were identified using MapMan software (v. 3.5.1R2, https://mapman.gabipd.org/home). The *Cannabis sativa* cs10 proteome was mapped to the MapMan-curated Arabidopsis TAIR10 pathways (downloaded 2020-March 10 from https://mapman.gabipd.org/mapmanstore) via BLASTP [57], filtering for alignments of cs10 – to −TAIR10 protein with percent ID >30% and >=50% of the shorter amino acid sequence length of the aligned pair.

## Results

### Microscopy

Cannabis capitate stalked glandular trichomes can be divided, morphologically, into two very distinct regions: the glandular trichome head and the stalk. The head appears as a bulbous sack on top of the stalk, connected by a short constricted neck that is continuous with the stalk **Fig 1[A]**. The base of the head is made up by a cluster of stipe cells that subtend the discoid layer of disc cells **Fig 1[B],**which are connected to the stalk. While the stipe cells emit red autofluorescence, indicative of photosynthetically active chloroplasts, the layer of dics cells are devoid of red autofluoresence, suggesting the absence of chloroplasts. The glandular trichome stalk is multicellular and is continuous with the epidermal layer **Fig 1[C]**. The cell boundaries of the glandular trichome stalk with the epidermis cannot be easily observed and indicate that the outer layer of stalk cells are similar in morphology to epidermal cells and develop perpendicular to the plane of epidermal cells. Fluorescence microscopy shows glandular trichome heads emit characteristic blue fluorescence indicative of the presence of phenolic compounds as previously observed in glandular trichomes from tomato [58], while glandular trichome stalks do not show this fluorescence **Fig 1[D]**. In contrast, chlorophyll autofluorescence (400-440nm excitation) was observed in glandular trichome stalks, suggesting stalk cells are photosynthetically active **Fig 1[B,D]**. In order to gain more insight into the unique function of the different trichome tissues a proteomic approach was undertaken.

**Figure 1.**
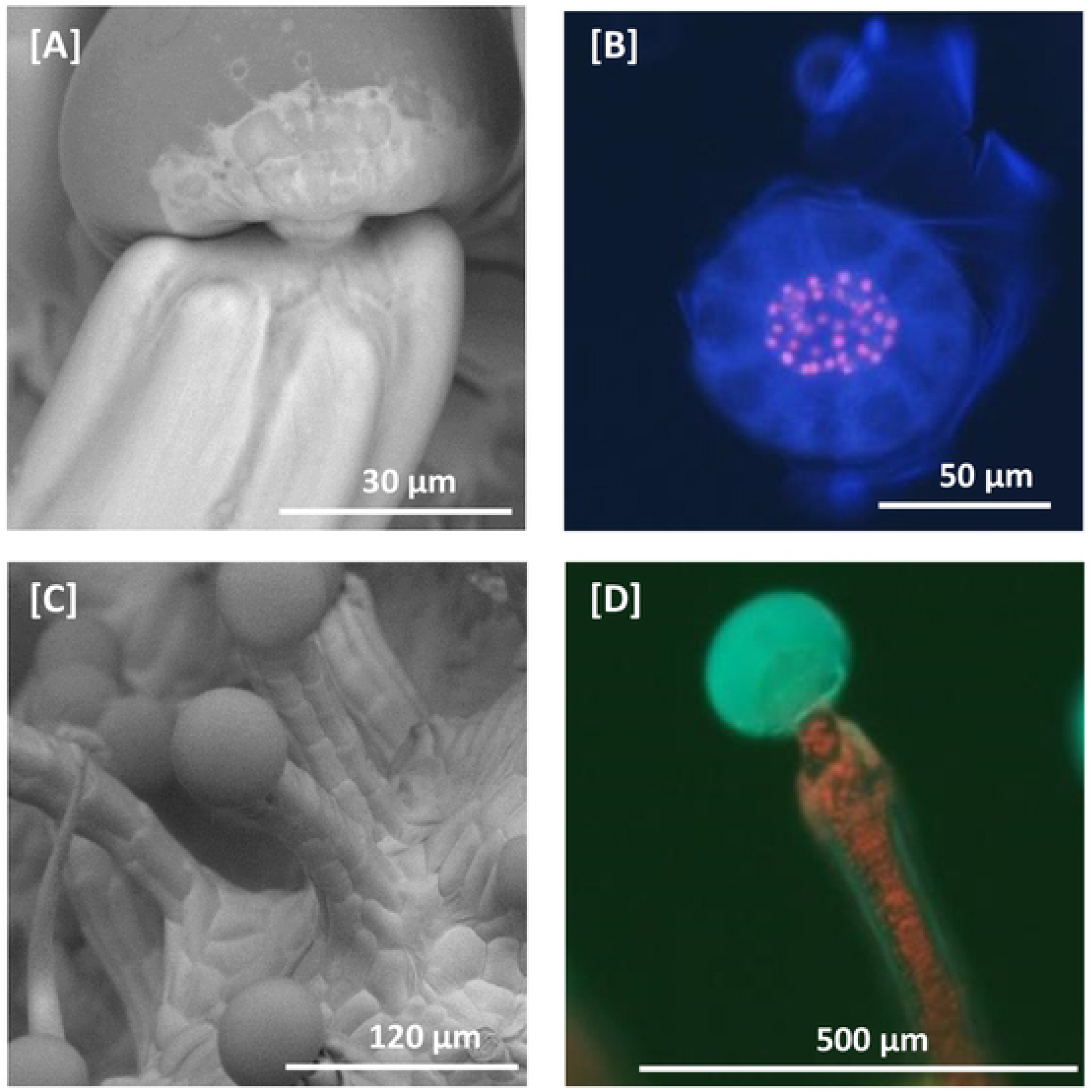
Scanning electron microscopy and fluorescent microscopy of *C. sativa* trichomes SEM micrograph of a capitate-stalked glandular trichome. Glandular trichome heads are morphologically distinct from the associated multicellular stalk. Glandular trichome heads are attached to multi-cellular stalks by a narrow constriction of the stalk. **[B]** Fluorescent microscopy image of a glandular trichome head showing the stipe cells (red fluorescence) that subtend the discoid layer of disc cells (no red fluorescence) at 400-440 nm excitation. Remnants of the ruptured cuticular layer covering the glandular cavity are visible on the right and top. Chlorophyll autofluorescence in red **[C]** SEM micrograph of capitate-stalked glandular trichomes, showing apparent continuity between epidermal layer and stalk. **[D]** Fluorescent microscopy image of capitate-stalked glandular trichome at 400-440 nm excitation with gland intact. Secondary metabolites in the glandular trichome head fluoresce blue, when excited, while chlorophyll in chloroplasts fluoresces red.

### Enrichment of trichome fractions

Glandular trichome heads and stalks were purified by Percoll density gradient centrifugation, from a heterogenous mix of epidermal tissues from excised flowers **Fig 2[A]**. Light microscopy confirmed that glandular trichome heads were recovered from the interface between 0% and 30% Percoll layers **Fig 2[B]**. Glandular trichome stalks were recovered from the interface between the 30% and the 45% Percoll layer **Fig 2[C]**. Tissues found at the 45%-55% and 55%-80% Percoll interfaces were comprised of mesophyll tissues and dense aggregates of heads and stalks and were excluded due to lack of purity (data not shown). Fragments of stigma and hair-type cystolithic trichomes were identified in the pellet following centrifugation. Although the glandular trichome head and stalk fractions were not entirely free from contamination i.e. some stalks were present in the glandular trichome head fraction **Fig 2[B]** and some heads were found in the glandular trichome stalk fraction **Fig 2[C]**there was considerable enrichment in the respective fractions to use them for downstream comparative proteome analysis between these tissue types.

**Figure 2.**
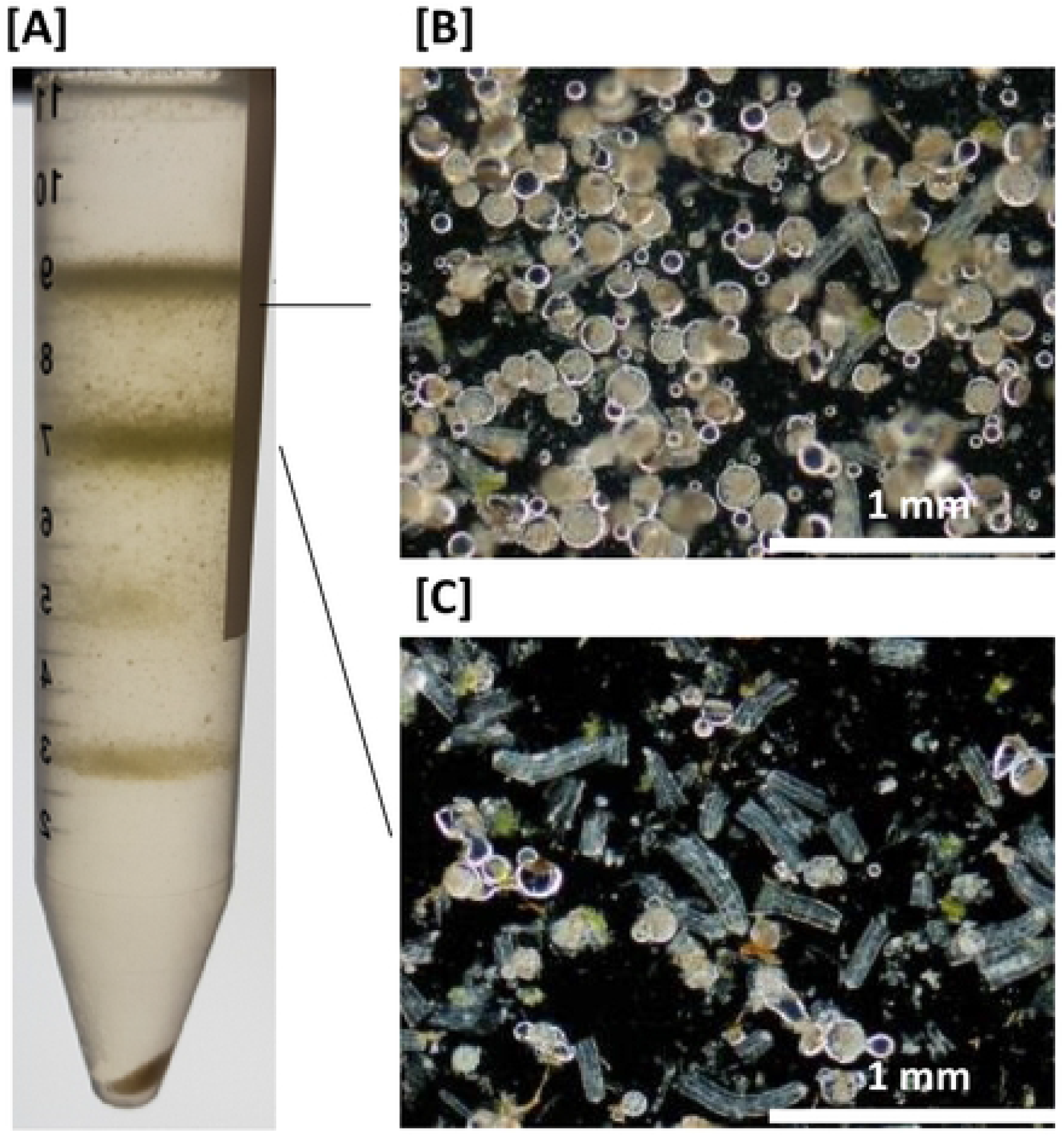
Purification of glandular trichome heads and glandular trichome stalks from a heterogenous mix of *C. sativa* flower epidermal tissues **[A]** Heterogenous mix of epidermal tissues separated by percoll gradient centrifugation. Glandular trichome heads collected at the interphase of the 0% and 30% Percoll boundary **[B]** and glandular trichome stalks collected at the interphase of the 30% and 45% Percoll boundary **[C]**

### Proteomics

Mass spectrometric data was searched against the cs10 *Cannabis sativa* reference proteome. The results obtained were analysed with Protein Pilot 5.0.2 software (SCIEX), then visualised and validated using Scaffold 4.8.7 (Proteome Science) employing a minimum protein threshold of 99.9% and peptide threshold of 95%. A total of 128,242 spectra were mapped to 1820 unique proteins across all tissues and replicates **Supplemental 1**. There were marked differences in the number of proteins identified between the sample types with 1682 proteins identified in late-stage *Cannabis sativa* flower, 1240 in glandular trichome heads, and 396 proteins were identified in glandular trichome stalks. Of the 1820 total proteins 569 (31.3%) were exclusive to late-stage flowers, while 122 (6.7%) were exclusive to glandular trichome heads and one protein (0.1%) was found to be unique to glandular trichome stalks **Fig 3[A]**.While 370 (20%) were found within all three sample types tested, 15 proteins (0.8%) were found in both glandular trichome heads and glandular trichome stalks, 10 proteins (0.6%) were found in glandular trichome stalks and late-stage flowers and 733 proteins (40%) found in glandular trichome heads and late-stage flowers **Supplemental Figure 1 [1-9]**.

**Figure 3.**
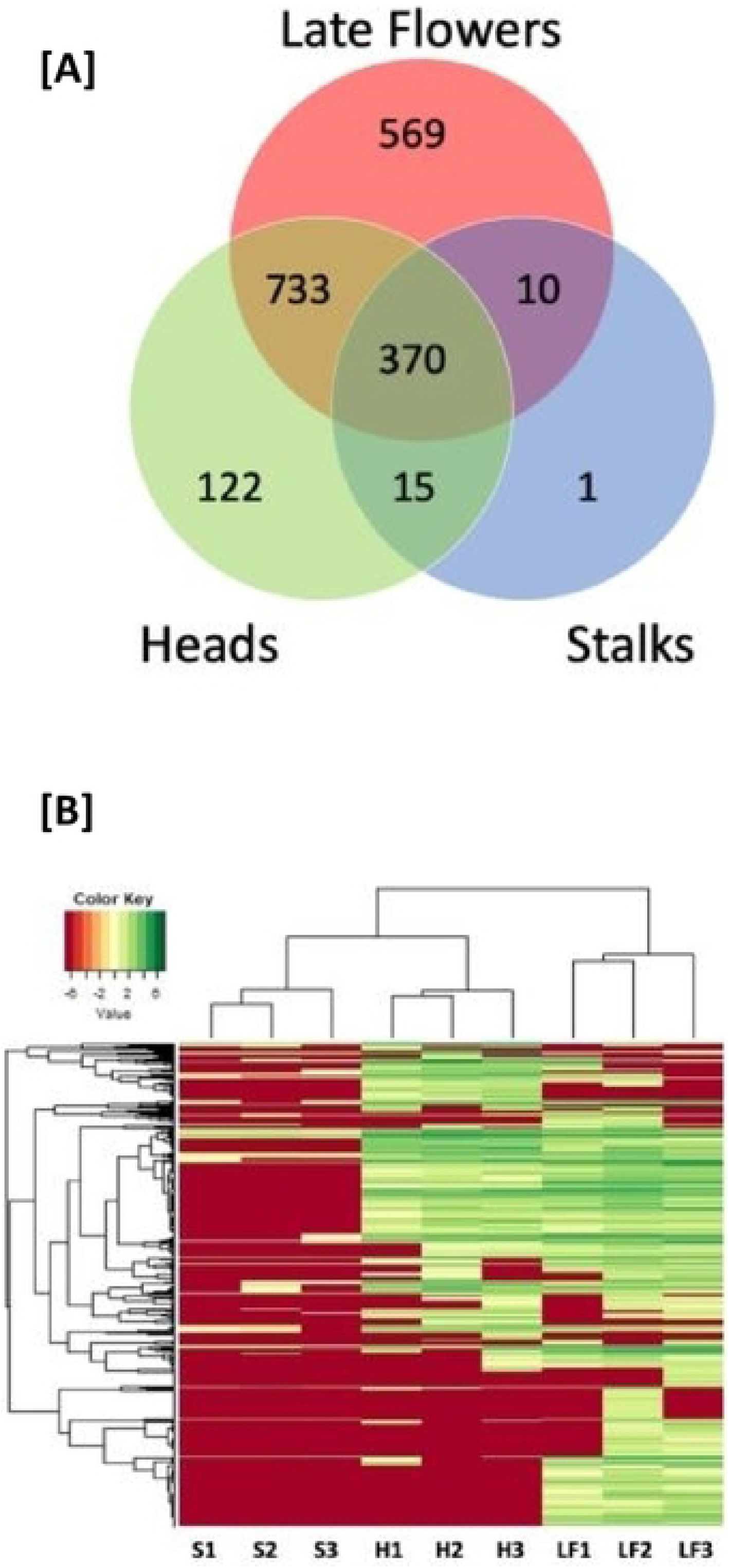
**[A]** Venn diagram comparing the proteins found within the three tissue types: glandular trichome heads, stalks, and late-stage *Cannabis sativa* flowers. **[B]** Hierarchical clustering of stalk enriched (S1-3), glandular trichome head (H1-3), and late flower (LF1-3) total spectral counts. The horizontal dendrogram represents the relationship each sample or column has with any other sample in the data set as pertains to that samples combination of spectral signatures. The vertical dendrogram defines the relationship a given protein or row has with any other protein in the data set with respect to abundance across all samples in this dataset.

Hierarchical clustering of total spectral counts across three biological replicates for each tissue type was applied to investigate the integrity of the data. Biological replicates for all samples were found to cluster together suggesting the three tissue types are distinct with respect to their proteomes. Furthermore, the glandular trichome head and stalk proteomes are more closely related to one another than either are to the late-stage flower proteome as indicated by the horizontal dendrogram **Fig 3[B]**.

Pairwise comparison of relative protein abundance between the three tissue types **Fig 4[A-C]**showed significant differences in protein abundance (blue circles) for individual proteins including presence/absence variation (red triangles). Of those proteins found in glandular trichome heads 4.2% were more abundant (p<0.0263; log2 fold change ≥ 1) when compared to late stage flowers while 2.4% of proteins found in late stage flowers were more abundant compared to heads (p <0.0263; log2 fold change ≥ 1) **Fig 4[A]**. 4.5% of proteins found in glandular trichome stalks were more abundant (p<0.02014 log2 fold change ≥ 1) compared to late-stage flowers while 19.2% of proteins found in late-stage flowers were more abundant (p<0.02014; log2 fold change ≥ 1) when compared to stalks **Fig 4[B]**. 14.3% of proteins found in glandular trichome heads were more abundant in heads compared to glandular trichome stalks (p<0.01009; log2 fold change ≥ 1), while 2.5% of proteins found in glandular trichome stalks were more abundant when compared to glandular trichome heads (p<0.01009; log2 fold change ≥ 1) **Fig 4[C]**. The majority of data points did not not pass the significance threshold (grey circles and triangles in **Fig 4[A-C]**).

**Figure 4.**
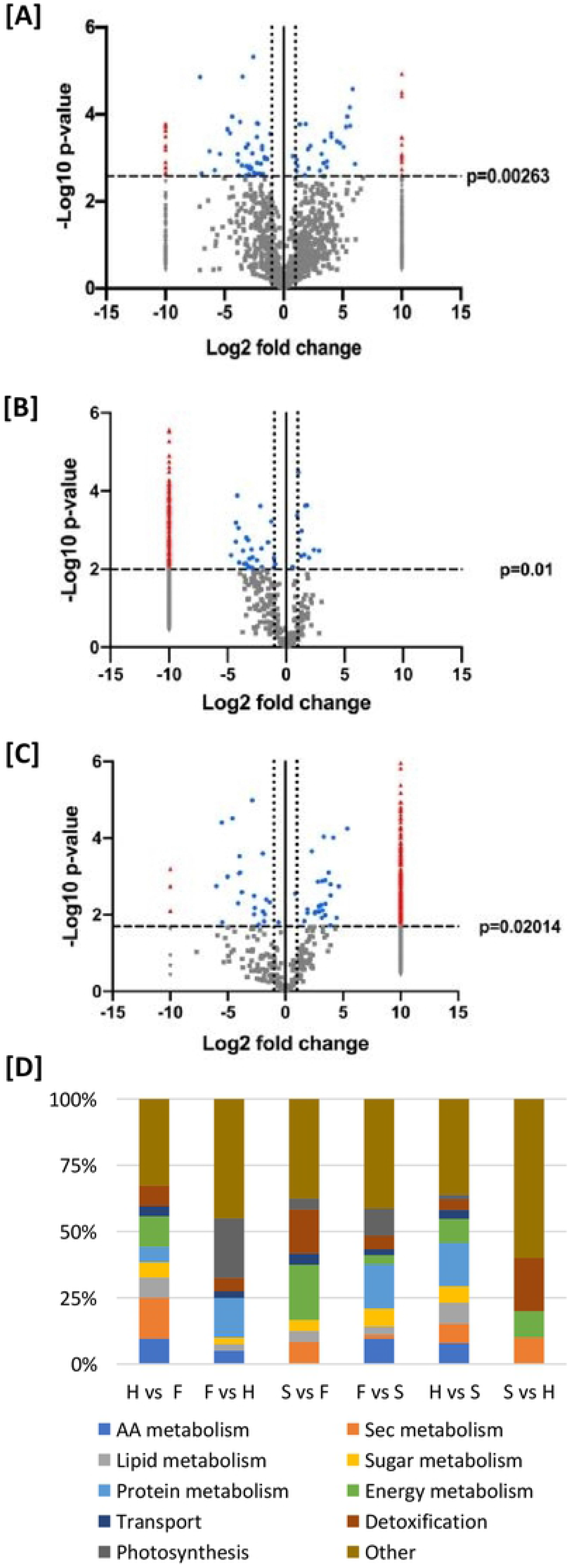
Volcano plot data models of relative protein abundance in a pairwise comparison of **[A]** glandular trichome heads and late-stage flowers. **[B]** Glandular trichome stalk enriched and late-stage flowers. **[C]** glandular trichome heads and glandular trichome stalk enriched. Each point on the model represents one protein. The Log2 (fold change: Heads/Late flower) is plotted against the –Log_10_(Benjamini-Hochberg corrected p-value). The vertical line at x=0 represents the 0-fold change threshold. Perforated lines intersecting x=−1 and x=1 represent a two-fold change in abundance. Red triangles denote proteins found exclusively in one group but not the other p<0.00263. Grey triangles represent proteins unique to one group that may be by chance. Blue dots represent proteins found in both sample types, that are significantly more abundant in one sample or the other. Grey dots represent proteins found in both sample types that are not significantly more abundant in either sample type. D: percent representation of major metabolic categories in pairwise comparisons …

### Comparison of Trichome Heads vs Flowers

Fifty-two proteins **Supplemental 2[1]** **Fig 4[A]** showed increased abundance in glandular trichome heads. Twelve were found to be mitochondrial or plastidal in origin. The largest group among these 52 proteins was related to secondary metabolism (15%), followed by energy metabolism (12%), amino acid metabolism (10%), lipid metabolism (8%) and detoxification (8%) **Fig 4[D]**. More than half of the proteins within the secondary metabolism category related to terpenoid and cannabinoid biosynthesis, including enzymes in the MEP pathway; isopentyl diphosphate delta isomerase (IDI) (ARE72265.1), alloromadendrene synthase (ARE72260.1), prenyltransferase 4 – CBGAS (DAC76710.1), olivetolic acid cyclase (AFN42527.1) and prenyltransferase 3-CsPT3 (DAC76713.1). Other secondary metabolite biosynthesis related proteins included chalcone isomerase (AFN42529.1) involved in flavonoid biosynthesis and strictosidine synthase-like 10 (XP_030482574.1) involved in alkaloid biosynthesis [59].

Forty proteins were found with increased abundance (p<0.00263) in late stage flowers **Supplemental 2[2]** and many were chloroplast derived proteins. Proteins related to photosynthesis were most abundant (23%), followed by protein metabolism (18%), detoxification (5%) and amino acid metabolism (5%) **Fig 4[D]**. Proteins related to photosynthesis included photosystem I (XP_030491785.1) and photosystem II (XP_030496367.1, XP_030482568.1) components as well as Calvin cycle proteins, while protein metabolism included translation related proteins, chaperones and disulphide isomerases. Other noteworthy proteins included elongation factors, and those related to chromatin assembly, detoxification, carotenoid degradation/pigmentation - carotenoid 9,10(9′,10′)-cleavage dioxygenase 1-like (XP_030493365.1) and fruit ripening-2-methylene-furan-3-one reductase (XP_030483081.1) **Supplemental 2[2]**.

Mapping of the highly abundant proteins in heads vs. late-stage flowers **Fig 5** indicated that late-stage flowers were comparatively enriched (blue colours) in photosynthesis and starch production, redox activity, and nitrogen assimilation while glandular trichome heads were comparatively more abundant in terpene and lipid biosynthetic processes as well as elements of the mitochondrial electron transport chain, citrate metabolism, and amino acid metabolism. These results indicated a primary sugar producing role for late-stage flowers while glandular trichome heads were strongly associated with the production and consumption of energy to drive secondary metabolite biosynthesis.

**Figure 5.**
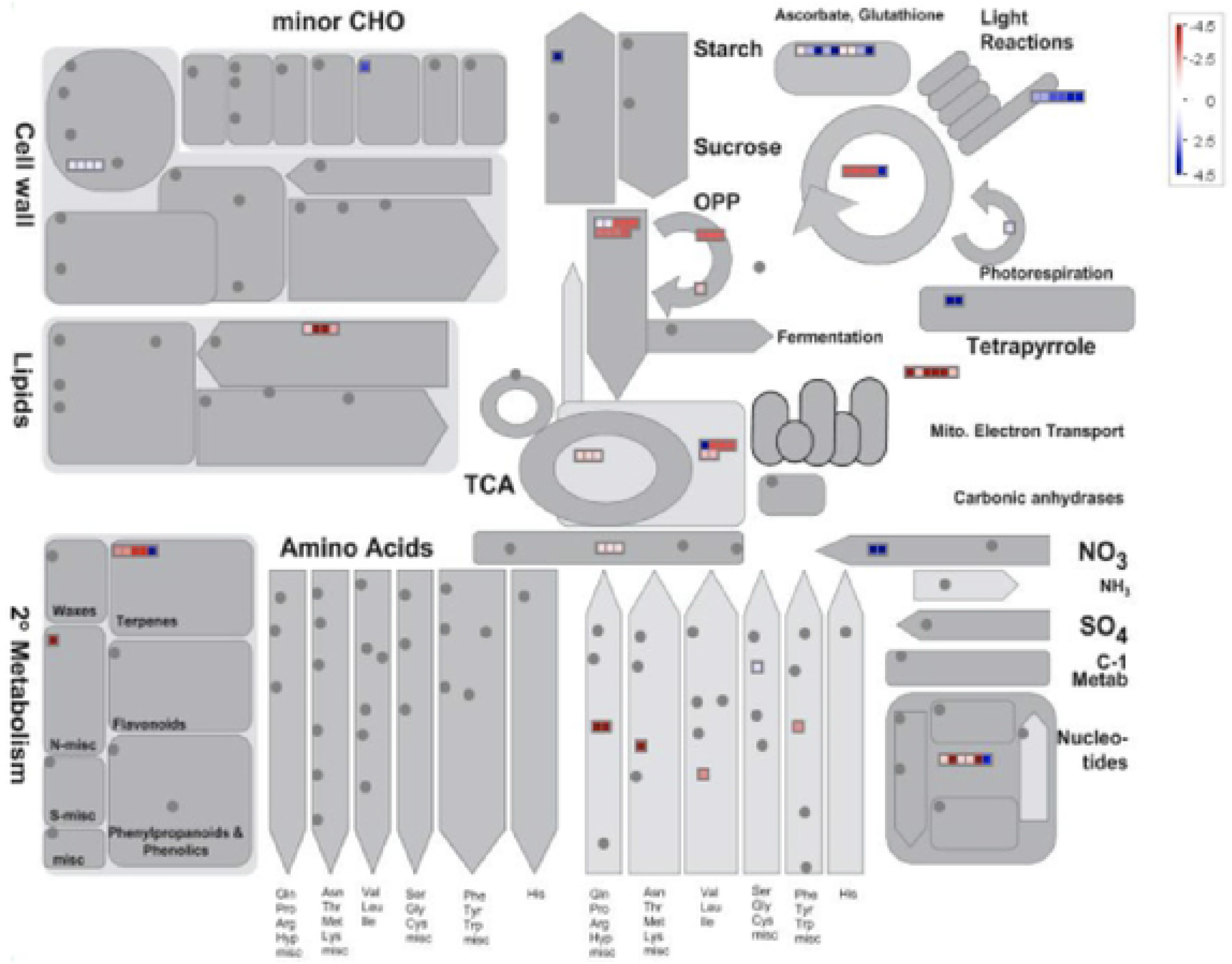
Mapman batch classification of proteins relating to metabolism in trichome Heads vs Flowers. Fold change differences in a protein’s abundance is indicated by the legend key where blue represents proteins found more abundant in late-stage flowers, white indicates no change in a protein’s abundance, and red indicates proteins found preferentially abundant in glandular trichome heads.

### Comparison of Trichome Stalks vs Flowers

Pairwise comparison of relative protein abundance between glandular trichome stalks and late-stage flowers is shown in **Fig 4[B]**. A total of 24 proteins were found to be increased in abundance (p <0.02014) in glandular trichome stalks **Supplemental 2[3]**. These proteins were mostly related to energy metabolism (20%) and detoxification (17%) **Fig 4[D]**. The remaining proteins included those involved in the MEP pathway for isoprenoid biosynthesis-HDS partial (ARE72266.1), cannabinoid biosynthesis-olivetolic acid cyclase (AFN42527.1) and cannabidiolic acid synthase (AKC34410.1), the citric acid cycle malate dehydrogenase (XP_030508797.1), chromatin assembly and transmembrane secondary active transport V-type proton ATPase subunit F (XP_030502093.1).

Three-hundered and fifty two proteins were found in increased abundance (p<0.02014) in late-stage flowers when compared to glandular trichome stalk proteins **Supplemental 2[4]**. Of those proteins 27% (97) were chloroplast derived proteins, and 9% (33) mitochondria derived. The primary GO categories of these more abundant proteins in late-stage flowers related to protein metabolism (17%), photosynthesis (10%) and amino acid metabolism (10%) **Fig 4[D]**.

Mapping of the proteins found more abundant in the pairwise comparison of glandular trichome stalks and late-stage flowers **Fig 6** indicated that many processes are largely underrepresented in the stalks when compared to the late-stage flower proteome. Notable exceptions to this include mitochondrial electron transport related proteins which were found to be more abundant in the stalks, while processes relating to nucleotide metabolism and terpene secondary metabolism were only modestly more abundant in the stalks. Late-stage flowers were comparatively more abundant in processes such as sugar and starch metabolism, cell wall biosynthesis, photosynthesis and respiration, and the biosynthesis of amino acids and lipids. The results indicated that glandular trichome stalks are limited in respect to their biological processes and most likely have a highly specialised biological role.

**Figure 6.**
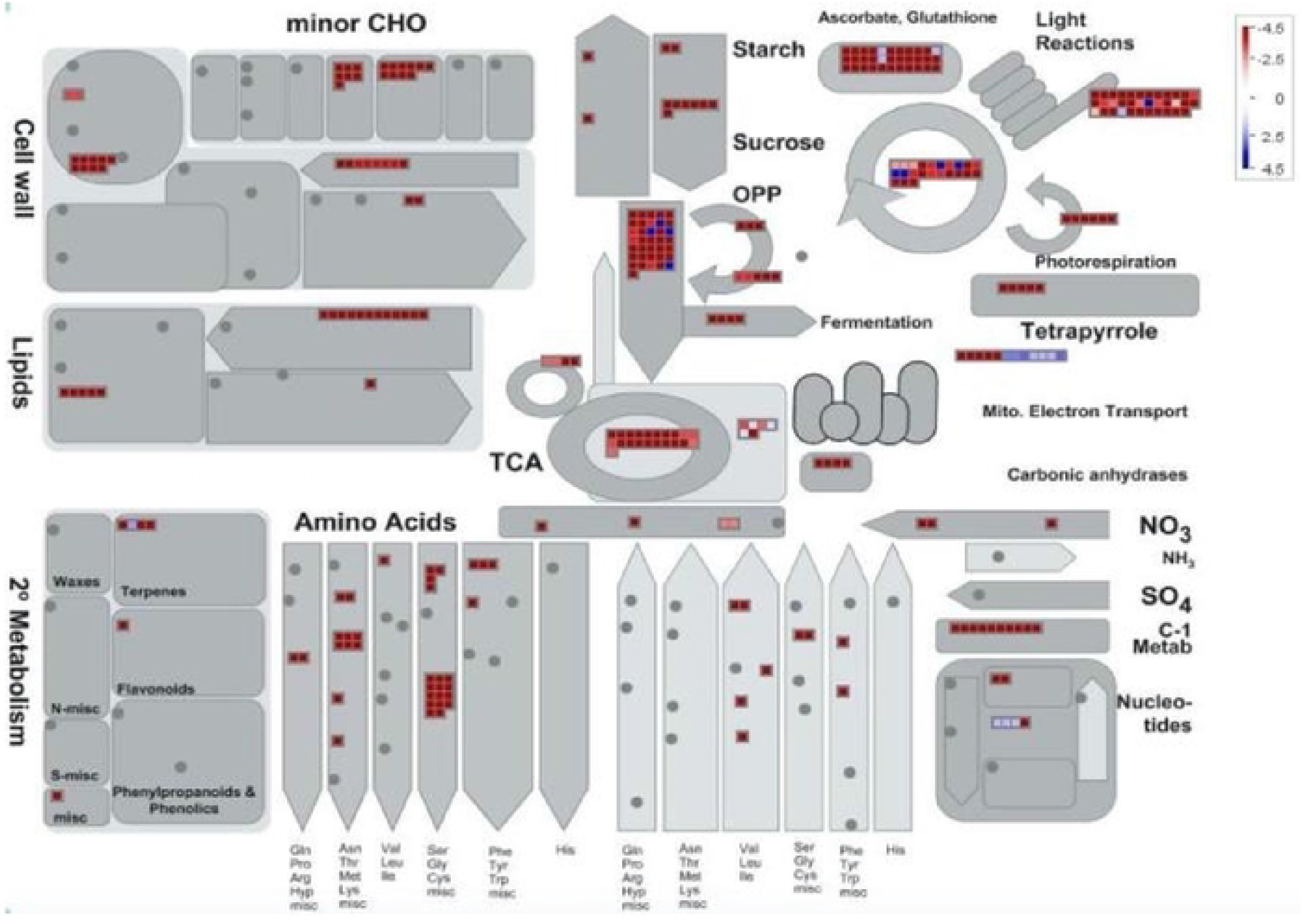
Mapman batch classification of proteins relating to metabolism in trichome Stalks vs Flowers. Fold change differences in a protein’s abundance is indicated by the legend key where blue represents proteins found more abundant in glandular trichome stalks, white indicates no change in a protein’s abundance, and red indicates proteins found preferentially abundant in late-stage flowers.

### Comparison of Trichome Heads vs Trichome Stalks

Pairwise comparisons of relative protein abundance between glandular trichome heads and glandular trichome stalks is shown in **Fig 4[C].** One hundred and seventy seven proteins displayed increased abundance in glandular trichome heads **Supplemental 2[5]**. These proteins were categorized into protein metabolism (16%) and energy metabolism (9%), followed by lipid metabolism (8%), amino acid metabolism (8%) and secondary metabolism (7%) **Fig 4[D]**.

Protein metabolism related proteins included: translation −60S ribosomal protein L10a (XP_030483002.1) and 40S ribosomal protein S19-3 like (XP_030484107.1), protein turnover - (ubiquitin-like protein SMT3 (CAI11094.1) and 26S proteasome non-ATPase regulatory subunit 4 homolog (XP_030485940.1). Energy metabolism included the citric acid cycle isocitrate dehydrogenase [NAD] regulatory subunit 1 (XP_030485874.1) and the electron transport chain-complex I (XP_030490934.1), complex III (XP_030500833.1), complex IV (XP_030496363.1), and ATP synthase subunit O (XP_030493792.1). Secondary metabolism included: MEP pathway -DXR (ARE72264.1), MCT (XP_030499547.1), and DXS (XP_030507676.1), terpene biosynthesis - alloaromadendrene synthase (ARE72260.1) and α-pinene synthase (ARE72261.1), cannabinoid biosynthesis – CBGAS prenyltransferase 4 (DAC76710.1), cannaflavin biosynthesis - CsPT3 (DAC76713.1), flavonoid biosynthesis - chalcone isomerase (AFN42529.1) and alkaloid biosynthesis - strictosidine synthase-like 10 (XP_030482574.1). Lipid metabolism included many proteins related to fatty acid biosynthesis related enzymes of particular interest was lineolate 13S lipoxygenase 2 (XP_030504574.1), a key enzyme in the biosynthesis of fatty alkyl groups, used in the biosynthesis of alkylresorcinolic acids - precursors to cannabinoid biosynthesis **Supplemental 2[5]**

Ten proteins were increased in abundance in glandular trichome stalks **Supplemental 2[6]** and were categorised into pathways/function **Fig 4[D].** Proteins with increased abundance in glandular trichome stalks (p<0.01) were associated with detoxification - peroxiredoxin-2E-2 (XP_030493857.1) and superoxide dismutase (XP_030504009.1), energy metabolism - ATP synthase F1 subunit (ALF004039.1) and secondary metabolism - olivetolic acid cyclase (AFN42527.1). Others included chromatin assembly – Histones: H2B.1 (XP_030504034.1), H4 (XP_030481652.1), and H2A.5 (XP_030492994.1) and, putative desiccation resistance - BURP domain protein RD22 (XP_030485614.1) and desiccation-related protein PCC13-62-like (XP_030492994.1).

Mapping of the proteins found increased in abundance in the pairwise comparison of both glandular trichome heads and glandular trichome stalks **Fig 7** indicated that glandular trichome heads (proteins shown in red) are hubs of secondary metabolic processes such as terpene, phenylpropanoid, and phenolic biosynthesis. Aside from secondary metabolism, glandular trichome heads were comparatively more abundant in ascorbate and glutathione metabolism, amino acid biosynthesis, sulfur metabolism, energy production and consumption. Glandular trichome stalks were comparatively lacking in notable biosynthetic processes, similar to the comparisson between glandular trichome stalks and late-stage flowers these results seem to indicate that glandular trichome stalks are likely highly specialised cells that are limited in the number of biological processes that they carry out.

**Figure 7.**
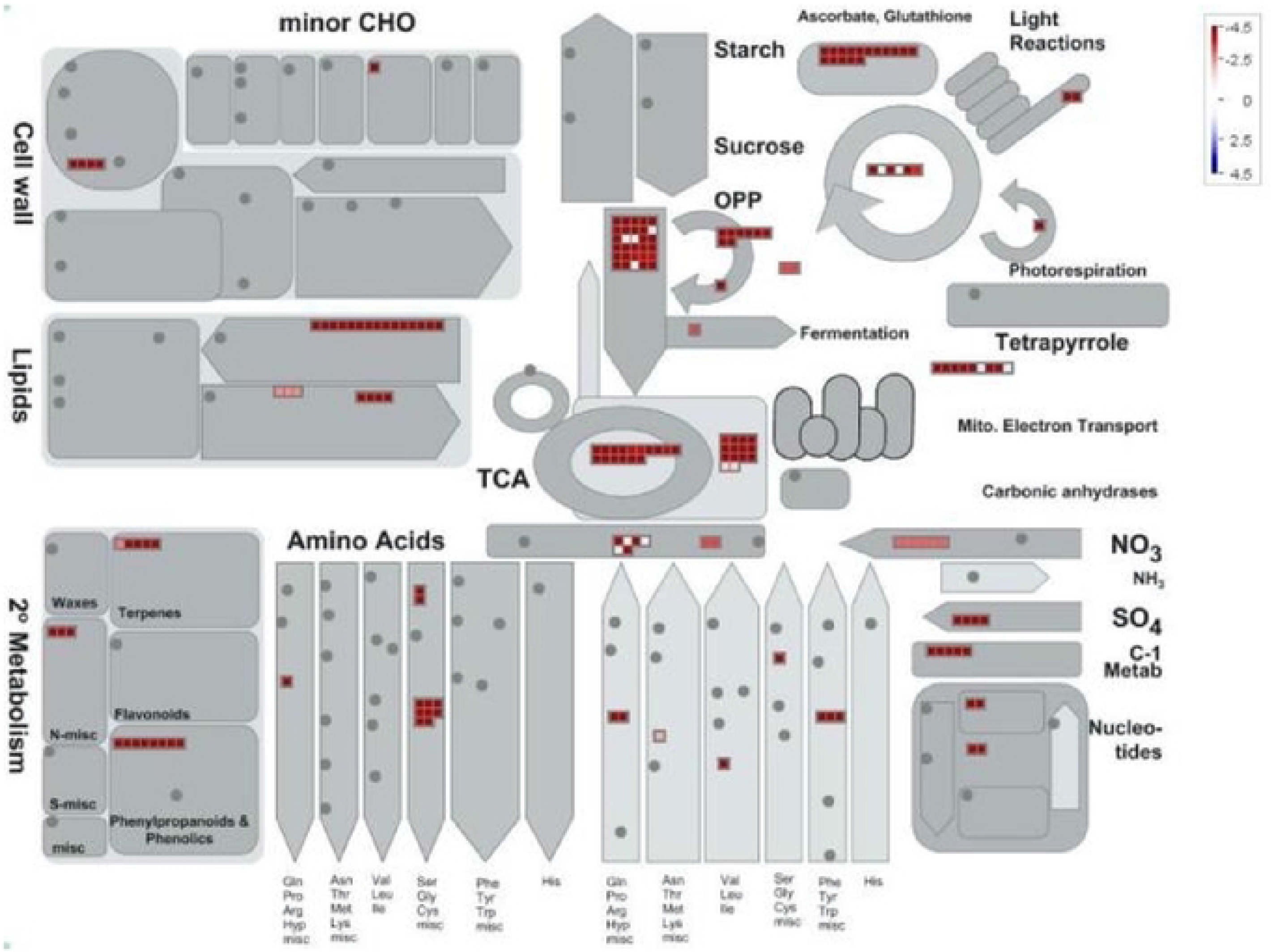
Mapman batch classification of proteins relating to metabolism in glandular trichome Heads vs Stalks. Fold change differences in a protein’s abundance is indicated by the legend key where blue represents proteins found more abundant in glandular trichome stalks, white indicates no change in a protein’s abundance, and red indicates proteins found preferentially abundant in glandular trichome heads.

## Discussion

Considering that major cannabinoids stored in trichome heads can account for over 15% of the *Cannabis sativa* female flower dry weight [60], trichome heads constitute a considerable carbon sink. In fact, in the absence of fertilization and subsequent grain filling, as is the case for medicinal *Cannabis* production, female floral trichomes can be assumed to be the major carbon sink during reproductive growth.

Though it cannot be ruled out that larger fan leaves contribute to source strength of *Cannabis sativa* during reproductive stages, it is more likely that female flowers themselves act as the primary source for carbon. While fan leaves showed clear signs of senescence, female inflorescences remained green and appeared healthy and photosynthetically active and are thus postulated to function as the main source tissue to fuel the filling of the trichome head subcuticular cavities during the late stages of flowering [61].

Pairwise comparisons of trichome heads, trichome stalks and whole late-stage flowers supported this model and provided a more detailed picture of source-sink interactions at the trichome level.

Around one quarter of the proteins displaying increased abundance in late-stage flowers compared to glandular trichome heads were related to photosynthesis and sugar production **Fig 4[D]** **&** **Fig 5**, indicating a strong role for late-stage flowers in the generation of photo-assimilates that is speculated to serve as a carbon and energy source for trichome head sink tissues. Source-sink-interactions and underlying metabolic signalling at reproductive stages during grain filling have been well studied [62–65]. Sucrose is a key signalling compound, which, through the activity of trehalose-6-phosphate (T6P) regulates primary metabolism. The signalling molecule T6P is thought to act as a sucrose gauge that has been demonstrated to directly inhibit SNF1 (sucrose-non-fermenting 1) related protein kinase (SnRK1) [66]. SnRK1 in turn acts as a central integrator of metabolic signals and governs over a switch from anabolism to catabolism in what is known as a feast or famine response. SnRK1 signalling cascades activate transcription factors that govern energy starvation responses [67]. Interestingly SNF-1 related protein kinase regulatory subunit gamma-1 like (XP_030495483.1) was found among the proteins in late-stage flowers **Supplemental 1[3]**. Presence of SnRK1 in late-stage flowers suggests that with the development of glandular trichomes a strong sink is established and that flow of carbon, likely in the form of C_6_ sugars, from flower mesophyll to trichome heads would be regulated through established metabolic signalling pathways.

Following a multi-omics approach to study glandular trichome function in tomato, sucrose has been proposed as a carbon and energy source for trichome sinks [41]. Tomato glandular trichomes however, are photosynthetically active, and for non-photosynthesising glandular trichomes, raffinose has been proposed as the C_6_ carbon source. In peppermint glandular trichomes for example raffinose has been demonstrated to fuel MEP-dependent terpenoid biosynthesis in leucoplasts [68].

In *Cannabis sativa,* chloroplast autofluorescence was present in trichome stalks **Fig 1[B,D]** and the stipe cells that subtend the disc cell layer of glandular trichome heads **Fig 1[B]**, but absent from the disc cells **Fig 1[B].** Distinction between chloroplast-containing stipe cells and chloroplast-devoid disc cells seems to suggest a functional separation of disc cells and stipe cells, which could not be further unravelled with the experimental set up. It is possible that the stipe cells are functionally more related to the cells of glandular trichome stalks i.e. playing a role in photosynthesis and sugar transport. Stipe cells are typically found with disc cells following separation of the glandular trichome heads from the glandular trichome stalk, **Fig 1[B]** which is not surprising considering they subtend the disc cell layer. The close relationship between the disc cells and stipe cells of *Cannabis sativa* glandular trichomes is shown by scanning and transmission electron microscopy studies [23, 69]. Several chloroplast-specific and photosynthesis associated proteins, were indeed found in the glandular trichome head fraction in this study, supposedly originating from stipe cells **Supplemental 1[1]** and is supportive of photosynthesis taking place in the heads. It has been proposed that light reactions in tomato photosynthetic trichomes supply the reducing power (NADH) demand of the MEP and MVA isoprenoid biosynthesis pathways and that carbon required for secondary metabolism is supplied from source tissues [41] or refixed respiratory CO_2_ [70]. The raffinose biosynthetic enzymes galactinol-sucrose galactosyltransferase 2 (XP_030493448.1) and 5 (XP_030482143.1) were detected in late-stage flowers **Supplemental 1[3]** while the enzyme alpha-galactosidase (XP_030508592.1), which catalyses the breakdown of raffinose into sucrose and D-galactose was found in glandular trichome heads **Supplemental 1[1]** suggesting a potential role for raffinose as a mobile photo-assimilate in *C. sativa* glandular trichome source-sink metabolism.

Moreover, in peppermint, a root specific isozyme ferredoxin NADP^+^ reductase, derived from non-photosynthetic tissues supplies electrons via a ferredoxin for the reductases of the MEP pathway, thus shunting production of monoterpenes via the MEP pathway enzymes (E)-4-Hydroxy-3-methyl-but-2-enyl pyrophosphate (HMB-PP) synthase (HDS) and HMB-PP reductase (HDR). In contrast, in photosynthetic glandular trichomes such as those from *Solanum habrochaites*, this non-photosynthetic ferredoxin reductase is missing, and terpenoids are derived from both the MEP and the MVA pathway [41, 68, 71]. Ferredoxin NADP reductase (XP_030479992.1) was indeed found in the glandular trichome head proteome **Supplemental 1[1]**, suggesting a high flux of MEP derived terpenoids in this cell type. This peppermint-like model for MEP pathway derived isoprenoids is further supported by the enrichment of MEP pathway proteins in glandular trichome heads **Supplemental 2[1]**, while MEV related proteins were present in both glandular trichome heads and late-stage flowers but not significantly more abundant in either. Interestingly, phosphoenolpyruvate (PEP)-carboxylase (PEPC) (XP_030480704.1), associated with CO_2_ fixation during C_4_ and CAM metabolism, showed increased abundance in heads when compared to both flowers and stalks **Supplemental 2[1&5]**. It has been speculated that PEPC could act in the refixation of respiratory CO_2_ in trichome heads to maximize carbon efficiency and supply for secondary metabolism [70, 72].

While our approach did not allow us to discriminate between the different cell types in the glandular heads, it suggested a model for *Cannabis sativa* glandular trichome secretory cells akin to the non-photosynthetic secretory system outlined by Shuurink & Tissier [73] - the layer of disc cells notably lacks chlorophyll autofluorescence **Fig 1[B]**, and is subtended by a layer of photosynthetically active stipe cells **Fig 1[B]**[23]. Enrichment of the oxidative pentose phosphate pathway **Fig [5 & 7]** as well as enrichment of the non-photosynthetic isozyme of ferredoxin reductase, which has been proposed to supply the reducing power to the enriched MEP pathway [68], shows similarity to non-photosynthetic secretory cells such as in peppermint, with specialised leucoplasts [74] as the site of secondary metabolite biosynthesis, relying on refixed carbon through PEP-carboxylase [70] and imported carbon in the form of sucrose/raffinose to maintain a high flux of MEP-derived terpenoids **Fig 8**.

**Figure 8.**
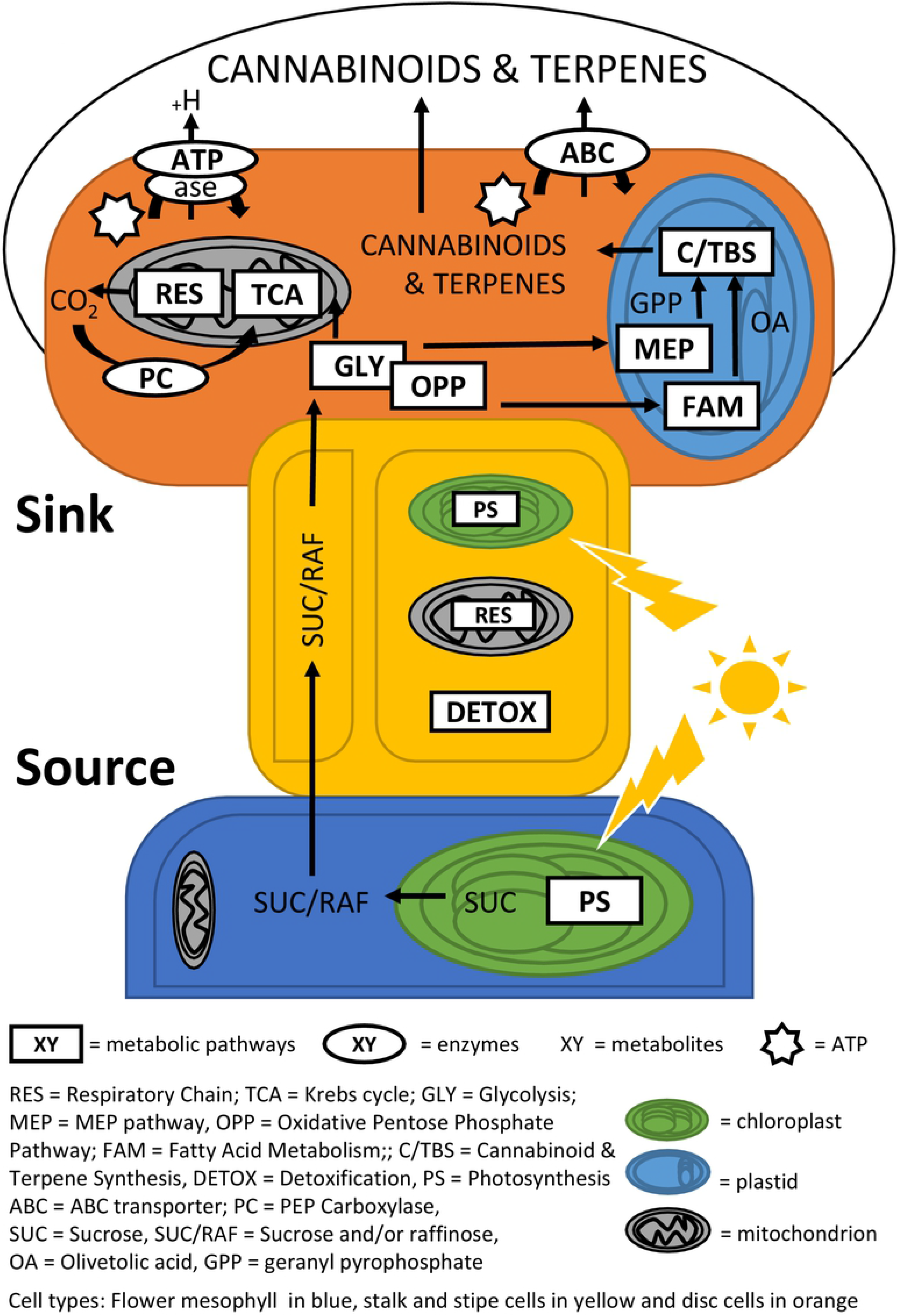
Carbon flow schematic of *Cannabis sativa* glandular trichomes, supported by proteomic data on late flower, trichome stalk and trichome head tissue. Mesophyll tissues of the late-stage flowers (blue) are source tissues for the photosynthetic (PS) production of sucrose (SUC). Sucrose and/or Raffinose (SUC/RAF) sugar is transported through glandular trichome stalks (yellow) to glandular trichome head sink tissue (orange) through yet unidentified mechanisms. Sugars are utilised by glandular trichome heads through glycolysis (GLY) and oxidative pentose phosphate pathway (OPP) to supply carbon to mitochondria and plastids. In mitochondria, catabolites enter Krebs cycle (TCA) and respiratory chain (RES) for energy production. Resulting carbon dioxide is partially recycled through pep-carboxylase (PC) activity. Resulting ATP is partially used to generate membrane gradients through ATPase activity and to drive primary active transport through ABC transporters (ABC). In plastids, catabolites can support terpenoid and cannabinoid biosynthesis (C/TBS) through the Methylerythritol Phosphate Pathway (MEP) and, in case of cannabinoids additionally through fatty acid metabolism (FAM). Stalk show PS and RES activity in addition to active detoxification (DETOX).

When compared to the late-stage flower proteome **Supplemental 2[3]**, stalk tissues showed an enrichment of the light dependent reaction enzyme photosystem II D1 (AJK91402.1), as well as chlorophyll autofluorescence throughout glandular trichome stalks **Fig 1[D]**. This suggested glandular trichome stalks play a role in light dependant photosynthesis. The glycolytic enzyme fructose bisphosphate 3-aldolase (XP_030496648.1), a marker for carbohydrate metabolism, including the Calvin cycle, was also highly abundant. We speculate, that the resulting sugars from stalk-based photosynthesis could act as an additional source of carbon and reducing power for glandular trichome heads or, equally likely, as local substrates to fuel high rates of mitochondrial respiration. In fact energy metabolism-related proteins (TCA cycle and respiratory chain components) made up the largest group of proteins with increased abundance in stalks (20%) compared to late flowers **Supplemental 2[3]**, followed by proteins relating to cellular detoxification (17%). High abundance of peroxidases and superoxide dismutase might indicate a harsh cellular environment rich in peroxides and other reactive oxygen species (ROS) in glandular trichome stalks. Peroxides and ROS abundance could be the result of high photorespiration and mitochondrial respiration rates respectively, supported by the observed enrichment of photosynthesis and mitochondria related proteins in glandular trichome stalks [75, 76]. In addition high irradiation under the controlled conditions of our experimental set up could also contribute to the production of superoxide species [77]. While disc cells are shielded from high irradiation by a large head space full of light absorbing terpenoids and cannabinoids, stalk cells are more exposed. Collectively, increased abundance of proteins involved in energy metabolism and detoxification in the stalks suggested a strong focus of these cells towards generation of ATP and reducing equivalents. ATP is speculated to be needed in stalk cells for membrane energization that drives primary or secondary active membrane transport. Subunits of the V-type proton ATPase that provide the electrochemical gradient to fuel secondary active transport [78], were found to be more abundant in the stalk proteome, compared with the late-stage flower proteome **Supplemental 2[3]**. The finding that they were not more abundant in the stalk proteome when compared with the head proteome might be due to the fact that membrane energization to maintain transport is also important in glandular trichome heads, which will be discussed below.

If C_6_ sugars are the mobile photo-assimilate in *Cannabis sativa* glandular trichomes, then the presence of secondary active C_6_ sugar transporters would be a reasonable assumption and could explain the need for high ATPase activity in the stalk cells. Plant sucrose symporters and antiporters are well described and characterized [79–81], however no sucrose or other related transporters were identified in the stalk proteome. Absence of any form of sugar transporters, could indicate that transport mechanisms other than secondary active trans-membrane transport are responsible for carbon shuttling from mesophyll through the stalk to the glandular trichome head. *Cannabis sativa* glandular trichomes are complex multicellular structures **Fig 1[A]** and there is a possibility that specialization within the stalk cells such as phloem-type vessels could facilitate the passive transport of sugar. Passive transport of sugars from mesophyll tissue via the plasmodesmata and loading phloem sieve tube elements is well documented in plants [82]. Research has shown that transport of small molecule markers does occur via plasmodesmata from epidermal cells to glandular trichome stalk cells in *N. tabacum* [83]. Interestingly, sieve element proteins (XP_030508649.1, XP_030508650.1, XP_030508648.1, and XP_030501215.1) were present in glandular trichome stalks and heads **Supplemental 1[1–2]** suggesting a model for *C. sativa* glandular trichome carbon supply through which sugar is transported from mesophyll source tissue to glandular trichome head sink tissue via sieve-like elements.

Surprisingly, two cannabinoid biosynthetic proteins (OAS, and CBDAS) displayed increased abundance in stalks vs the late-stage flower proteomes. While this could indicate contribution of the stalks to biosynthesis of cannabinoids, it could also be a result of contamination of the stalk fraction with disc cell proteins from the heads during the isolation via gradient centrifugation **Fig 2[C]**.

In conclusion, while there is a clear indication of Cannabis glandular trichomes stalks contributing to energy production, photosynthesis and detoxification, however, details on how glandular trichome stalks take part in active sugar transport and the biosynthesis of secondary metabolites remains unclear.

Taken together these results support a model for carbon generation, flow and utilization within glandular trichomes and underlying mesophyll tissue **Fig 8**. C_6_ sugars, likely raffinose or sucrose, are produced by the mesophyll cells of the late-stage flower and transported via yet unknown mechanisms in trichome stalks to stipe and disc cells at the base of the trichome heads. There, they would be broken down into simple sugars and enter sugar catabolic pathways including glycolysis (GLY) and the oxidative pentose phosphate pathway (OPP) [84]. Indeed, a total of 6% of overly abundant proteins in heads compared to stalk proteins and 4% in the comparison between heads and flowers were associated with these pathways **Supplemental 2[1&5].**

C3 pyruvate can enter energy metabolism, TCA cycle and respiratory chain (RES), for ATP production. Nine percent (9%) of the proteins with increased abundance in heads vs stalks are associated with energy metabolism, suggesting high rates of energy production. In the case of trichome heads compared to late stage flowers that number is 12% **Fig 4[D].**The comparatively higher demand for energy in glandular trichome heads supports an extensive secondary metabolism and possibly membrane energization **Supplemental 2[5]**. Intermediates and end products of glycolysis (GLY) and pentose phosphate pathway (OPP) can enter the fatty acid metabolism (FAM) pathway to support lipid biosynthesis needed to support the increase in membranes required for trichome head expansion and possible vesicular trafficking of secondary metabolites, as well as to provide precursors such as olivetolic acid for cannabinoid biosynthesis. Eight percent of the overly abundant proteins in heads vs stalks and 8% in heads vs late flowers are associated with lipid metabolism **Fig 4[D],**underlining the importance of these pathways in glandular trichome heads. Ultimately the majority of carbon and energy reaching the disc cells of trichome heads is speculated to be invested in secondary metabolite biosynthesis pathways including cannabinoid and terpenoid biosynthesis. While cannabinoid, terpenoid, and MEP pathway biosynthesis associated proteins make up 5 % of the proteins increased in abundance in the head vs stalk, they make up 10% in head vs flower. Proteins contributing to other secondary metabolite pathways make up 3% in head vs stalk and 6% in head vs flowers of the enriched proteins. Notably, 10% of proteins found preferentially more abundant in glandular trichome heads compared to late stage flowers were related to amino acid (AA) metabolism. The abundance of AA metabolism related proteins supports the hypothesis of glandular trichome heads as secondary metabolite biosynthetic factories, as AA’s serve as precursors for many of these compounds [85]. Moreover, P-type, V-type, and mitochondrial ATPases are found comparatively more abundant in heads compared to late-stage flowers **Supplemental 2[1]**, presumably to support the filling of subcuticular trichome cavities with secondary metabolites driven by proton (H^+^) gradients [78, 86], and to produce ATP to fuel the diverse primary and secondary metabolism of glandular trichome heads [87]. With respect to secondary metabolite transport, an ABC transporter B family member (XP_030500887.1) and pleiotropic drug resistance protein 1-like (XP_030488377.1) were found only in glandular trichome heads **Supplemental 1[1]**. Plant PDR and ABC subfamily B transporters are largely plasma membrane localised transporters that function in the efflux of secondary metabolites from cells via the hydrolysis of ATP [88], contributing to the overall demand for ATP. A PDR type transporter of petunia has previously been implicated in the trichome-specific transport of steroidal compounds or their precursors, contibuting to trichome-mediated herbivory defense [89].

Our model supports the paradigm of glandular trichomes as highly specialised secondary metabolite factories and contributes to the understanding of *Cannabis sativa* glandular trichome biology. By drawing on the differences and similarities in protein composition between the components of glandular trichomes and flowers we have developed an initial framework for the functions and processes that govern *Cannabis sativa* glandular trichome productivity. Our findings lay the foundation for future research into metabolic regulation of medicinally important secondary metabolites in *Cannabis sativa* glandular trichomes.

## Acknowledgements

The authors would like to acknowledge Alun Jones, Manager, Mass Spectrometry Facility, IMB University Queensland for MS analysis, Maxine Dawes, SCU, for technical support for SEM, Nicolas Dimopoulous for contribution of Fig 1[B], Andrew Kavasilas for donating the hemp cultivar used in this research, Michael Karkkainen for his input on matters relating to compliance and handling of controlled substances, Qi Guo for support with proteomic data handling and Priyakshee Borpatra Gohain and Francine Gloerfelt-Tarp for help with maintaining the plants. LJC was supported by a joint MSc scholarship supported by Cann Group Ltd and Southern Cross University.

